# ENDOSOMAL SIGNALING OF PROTEASE-ACTIVATED RECEPTOR-2 AMPLIFIES HISTAMINE-INDUCED PAIN OF IRRITABLE BOWEL SYNDROME

**DOI:** 10.64898/2026.01.30.702691

**Authors:** Nestor N Jimenez-Vargas, Badr Sokrat, Daniella A Gilmour, Maria F Pessano, Steve Davidson, Brian L. Schmidt, David E Reed, Alan E Lomax, Stephen J Vanner, Nigel W Bunnett

## Abstract

**Background:** Proteases and histamine, co-secreted by mast cells and bacteria, sensitize colonic nociceptors and contribute to irritable bowel syndrome (IBS) pain.

**Objective:** To determine whether irreversible proteolytic cleavage of protease-activated receptor-2 (PAR_2_) and its continued activity in endosomes amplify and sustain otherwise transient pronociceptive actions of histamine receptors (HRs) to cause recurrent pain, the defining symptom of IBS.

**Design:** We investigated the coexpression of PAR_2_ and H_1_R in nociceptors using RNAscope *in situ* hybridization and assessed the consequences of coactivation using electrophysiological assays of nociceptor sensitization and biophysical measurements of receptor and effector activity.

**Results:** PAR_2_ and H_1_R were coexpressed by human and mouse dorsal root ganglion nociceptors. Intracolonic infusion of fecal supernatants from IBS patients enhanced mechanosensitivity of colonic nociceptors in mice. Antagonists of PAR_2_ or H_1-4_R abolished this response. Combined administration of subthreshold concentrations of trypsin and histamine replicated the effects of fecal supernatant and caused hyperexcitability of isolated nociceptors. Pre-activation of PAR_2_ sensitized histamine-induced hyperexcitability. Endocytosis inhibitors prevented this hypersensitivity, consistent with sustained endosomal signaling of PAR_2_ and persistent nociceptor hyperexcitability. Trypsin amplified histamine-induced activation of H_1_R and β-arrestin2 and Gαq effectors at the plasmalemma and in endosomes. Conversely, histamine did not sensitize trypsin-induced hyperexcitability of neurons, in line with the inability of histamine to induce sustained nociceptor hypersensitivity.

**Conclusions:** By amplifying and maintaining the otherwise transient actions of H_1_R and possibly other pain receptors, persistent PAR_2_ endosomal signaling makes a dominant contribution to IBS-related colonic pain.

**Summary box:** *What is already known on this topic:* Proteases and histamine are increased in IBS patients and cause visceral pain.

*What this study adds:* Prolonged intracellular PAR_2_ signaling sensitizes and maintains H_1_R activity to amplify and maintain pain.

*How this might affect research, practice or policy:* Although neuroactive factors can act synergistically to amplify and maintain IBS pain, antagonists of dominant receptors (*e*.*g*., PAR_2_) can provide effective treatment.

## Introduction

Visceral pain is a hallmark of gut-brain interaction disorders, including IBS [1]. These disorders are common, debilitating, poorly understood and inadequately treated [2]. Proteases and histamine, released from infiltrating mucosal mast cells in IBS patients, sensitize colonic nociceptors [3, 4, 5, 6]. Tryptase and trypsin cleave PAR_2_ [7, 8] and histamine activates nociceptor H_1_R [6, 9], causing nociceptor hyperexcitability and visceral pain. Colonic bacteria also secrete proteases and histamine [10, 11, 12]. *Bacteroides fragilis* secretes the serine protease *B. fragilis* protease 1, which activates PAR_2_ on nociceptors to cause visceral pain [13]. *Klebsiella aerogenes* is a major source of histamine in fecal samples from IBS patients that can sensitize nociceptors [11].

PAR_2_ and H_1_R are G protein-coupled receptors (GPCRs) with distinct mechanisms of activation, signaling and regulation. Whereas histamine interacts reversibly with HRs, proteases irreversibly cleave PAR_2_ to reveal a tethered ligand that is always available to interact with the cleaved receptor to cause persistent activation [14]. GPCR kinases phosphorylate activated GPCRs at the plasma membrane, increasing their affinity for β-arrestins, which mediate receptor desensitization and endocytosis [15]. While these mechanisms terminate H_1_R signaling at the plasma membrane, PAR_2_ continues to signal from endosomes by G protein and β-arrestin-dependent mechanisms that sensitize nociceptors and contribute to IBS-associated visceral pain [16, 17, 18, 19]. Whether H_1_R signaling at the plasma membrane and PAR_2_ signaling in endosomes converge to sensitize nociceptors and trigger visceral pain is unknown.

The observations that proteases and histamine are upregulated in IBS patients and sensitize nociceptors suggest that PAR_2_ and H_1_R function cooperatively to induce visceral pain. Protease inhibitors and HR antagonists independently yet completely block the hyperexcitability of colonic afferent nerves in mice induced by IBS fecal supernatant (FS) [11, 12], which supports this proposal. However, despite the likelihood that proteases and histamine signal in concert, most studies have examined PAR_2_ and H_1_R signaling independently. We investigated the hypothesis that irreversible proteolytic activation of PAR_2_ and its propensity to generate persistent signals from endosomes amplify and sustain the otherwise transient pronociceptive actions of the H_1_R to cause recurrent pain, the defining feature of IBS.

## Results

### PAR_2_ and H_1-4_Rs mediate IBS FS-induced mechanical hypersensitivity of colonic nociceptors

To determine whether feces from IBS patients contains factors that cause hypersensitivity of colonic nociceptors, we administered FS or vehicle (Krebs solution) into the lumen of isolated segments of mouse distal colon and measured action potential firing in colonic afferent axons under basal conditions and in response to graded colonic distension. Fecal samples were obtained from patients that fulfilled the Rome IV criteria for diarrhea-predominant IBS (IBS-D). FS was prepared from 3 patients with high IBS symptom severity score (IBS-SSS, >300, **Table S1**) and high Patient-Reported Outcomes Measurement Information System (PROMIS) belly raw data pain scores (>19, **Tables S2, S3**). IBS-D FS significantly increased the basal firing frequency of colonic afferents compared to control (IBS-D FS, 0.65±0.12 Hz; control, 0.28±0.06 Hz; Wilcoxon paired test, *P*<0.001, N=15 mice, n=3 IBS-D FS) (**Fig. 1A**). IBS-D FS significantly increased responses to colonic distension at pressures >30 mm Hg, which are mediated by high-threshold fibers [20], compared to control (F (1, 13)=21.82 [treatment effect], 2-way RM ANOVA, *P*<0.001; 60 mm Hg: IBS-D FS, 148.7±16.5%; control, 92.9±2.9%; Šídák’s *post hoc* test, *P*<0.05). FS from healthy volunteers does not affect mechanosensitivity of colonic afferent neurons or excitability of nociceptors [12, 21].

**Figure 1.**
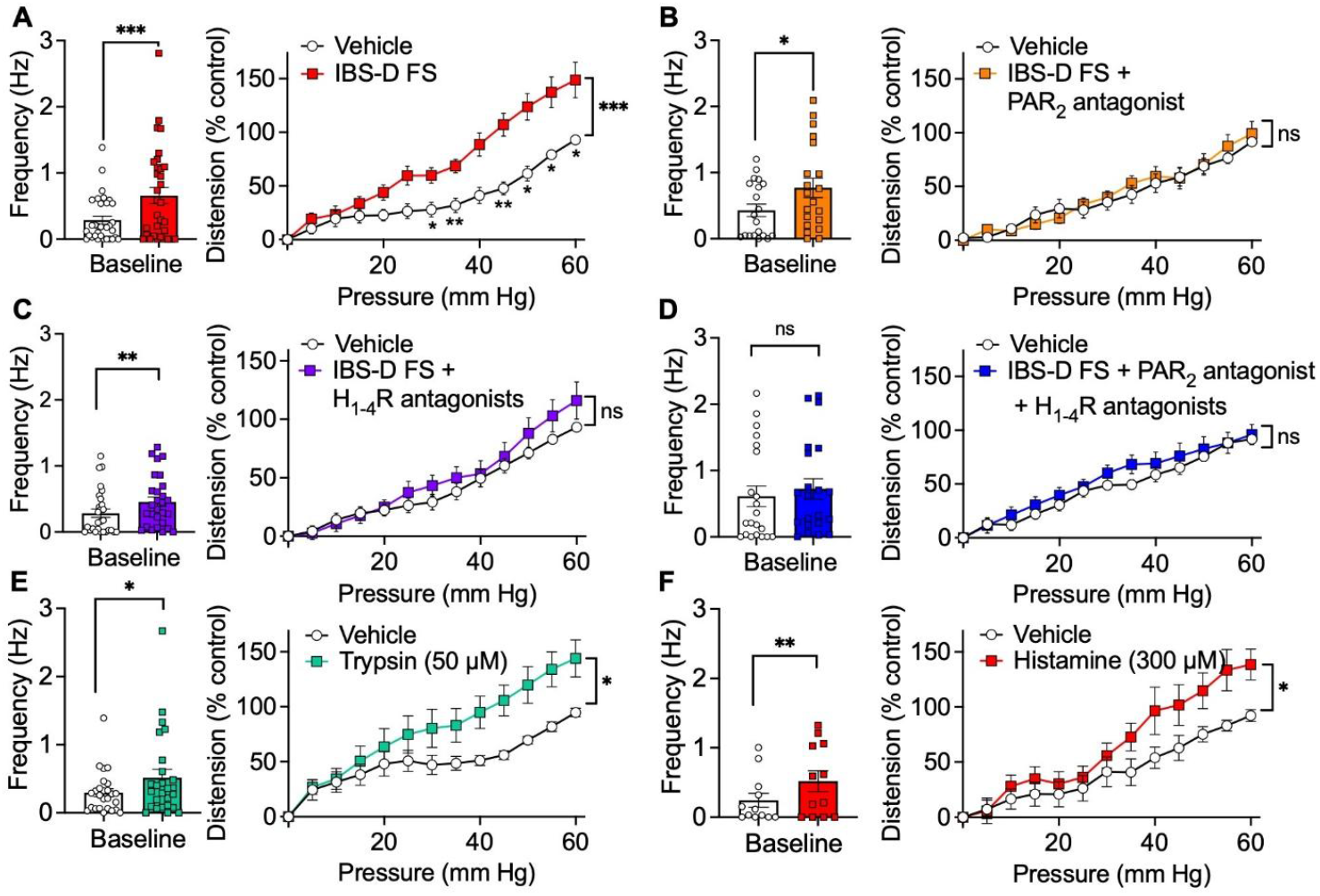
Activity of colonic afferent neurons after intraluminal administration of IBS-D FS, trypsin, histamine or vehicle. Baseline activity and distension-induced responses of colonic afferent neurons after intraluminal administration of IBS-D FS (**A-D**), trypsin (50 µM) (**E**), histamine (300 µM) (**F**) or vehicle (control). Tissues were pre-treated with vehicle (control) (A), PAR_2_ antagonist (GB83, 10 µM) (B), H_1-4_R antagonists (1 μM pyrilamine, 10 μM ranitidine, 30 nM clobenpropit, 1 μM JNJ7777120) (C), or a combination of PAR_2_ and H_1-4_R antagonists (D). Mean±SEM; N=number of mice, n=number of IBS-D FS patients, U=number of neuronal units. A: N=14, n=3, U=32. B: N=13, n=2, U=20. C: N=13, n=3, U=28. D: N=13, n=3, U=21. E: N=10, U=26. F: N=8, U=12. \**P*< 0.05, \*\**P*< 0.01, \*\*\**P*< 0.001, 2-way ANOVA with Šidák’s *post hoc* test or Wilcoxon paired test.

To determine the contribution of receptors for proteases and histamine to the IBS-D FS responses, tissues were pre-treated with antagonists of PAR_2_ (10 µM GB83) or H_1-4_Rs (H_1_R, 1 µM pyrilamine; H_2_R,10 µM ranitidine; H_3_R, 30 nM clobenpropit; H_4_R, 1 μM JNJ7777120). Antagonism of PAR_2_ or H_1-4_R did not affect the increased basal activity of colonic afferents to IBS-D FS but completely prevented response of IBS-D FS-treated preparations to colonic distension (**Fig. 1B, C**). PAR_2_ plus H_1-4_R antagonists inhibited both the basal and distension-induced activity of colonic afferents to IBS-D FS (**Fig. 1D**). PAR_2_ or H_1-4_Rs antagonists did not affect basal activity or distension responses of untreated colon (*i*.*e*., no FS) (**Fig. S1**). Thus, receptors for proteases and histamine independently mediate IBS-D FS-induced mechanical hypersensitivity of colonic afferents.

### Trypsin and histamine synergistically increase mechanosensitivity of colonic nociceptors

To determine whether selective activation of PAR_2_ and H_1-4_Rs replicate IBS-D FS-induced responses, trypsin (50 µM) or histamine (300 µM) were perfused through the colon lumen. Trypsin and histamine significantly increased basal activity of colonic afferents compared to control (trypsin, 0.51±0.12 Hz; control 0.29±0.05 Hz; Wilcoxon signed rank test, *P*<0.05; histamine, 0.51±0.15 Hz; control 0.24±0.1 Hz; Wilcoxon signed rank test, *P*<0.01) (**Fig. 1E, F**). Trypsin and histamine significantly increased responses to distension compared to control (trypsin: F (1, 9)=5.59 [treatment effect], N=10 mice; histamine, F (1, 7)=5.82 [treatment effect], N= 8 mice; 2-way RM ANOVA, *P*<0.05) (**Fig. 1 E, F**). These results align with reports that trypsin and histamine, respectively, activate PAR_2_ and HRs on nociceptors to evoke hyperexcitability [6, 22].

Subthreshold concentrations (*i*.*e*., that did not increase colonic afferent nerve mechanosensitivity compared to control during distension) of trypsin and histamine were determined experimentally to evaluate whether their co-application would increase colonic afferent mechanical sensitivity. Intracolonic trypsin (30 µM) or histamine (30 µM) did not significantly affect basal activity or mechanosensitivity of colonic afferents compared to vehicle (**Fig. 2A, B**). In contrast, co-application of subthreshold trypsin (30 µM) plus histamine (30 µM) significantly increased basal firing of colonic afferent nerves (co-application, 0.66±0.09 Hz; control 0.4±0.05 Hz; Wilcoxon signed rank test, *P*<0.001, N=18) and distension-induced responses at pressures above 40 mmHg (F (1, 17)=18.15 [treatment effect], 2-way RM ANOVA, *P*<0.001; 60 mm Hg: co-application, 141.8±11.4%, control, 95±1.3%; Šídák’s *post hoc* test, *P*<0.05) (**Fig. 3C**). Thus, co-application of concentrations of trypsin and histamine, which individually have no effect, increases colonic afferent nerve mechanosensitivity. These results suggest synergistic interactions between PAR_2_ and HRs contribute to the sensitization of colonic nociceptors to mechanical stimulation, a feature of IBS [1].

**Figure 2.**
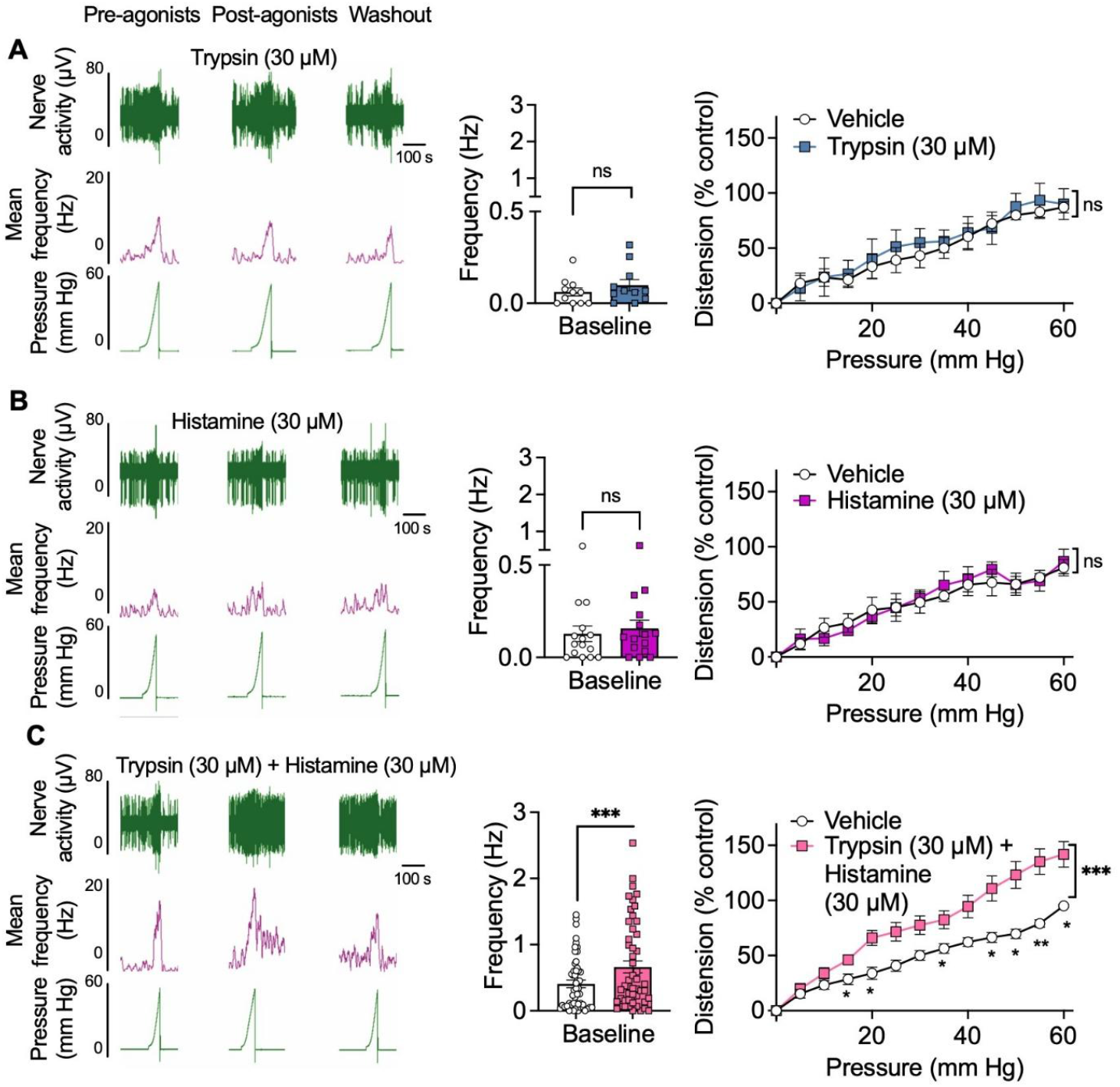
Activity of colonic afferent neurons after intraluminal administration of subthreshold concentrations of trypsin, histamine or vehicle. Representative traces, baseline activity and distension-induced responses of colonic afferent neurons after intraluminal administration of subthreshold concentrations of trypsin (30 µM) (**A**), histamine (30 µM) (**B**), both trypsin (30 µM) and histamine (30 µM) (**C**) or vehicle. Mean±SEM, N=number of mice, U=number of neuronal units. A: N=6, U=11; B: N=6, U=15. C: N=18, U=51 \*\*\**P*< 0.001, 2-way ANOVA with Šidák’s *post hoc* test or Wilcoxon paired test.

**Figure 3.**
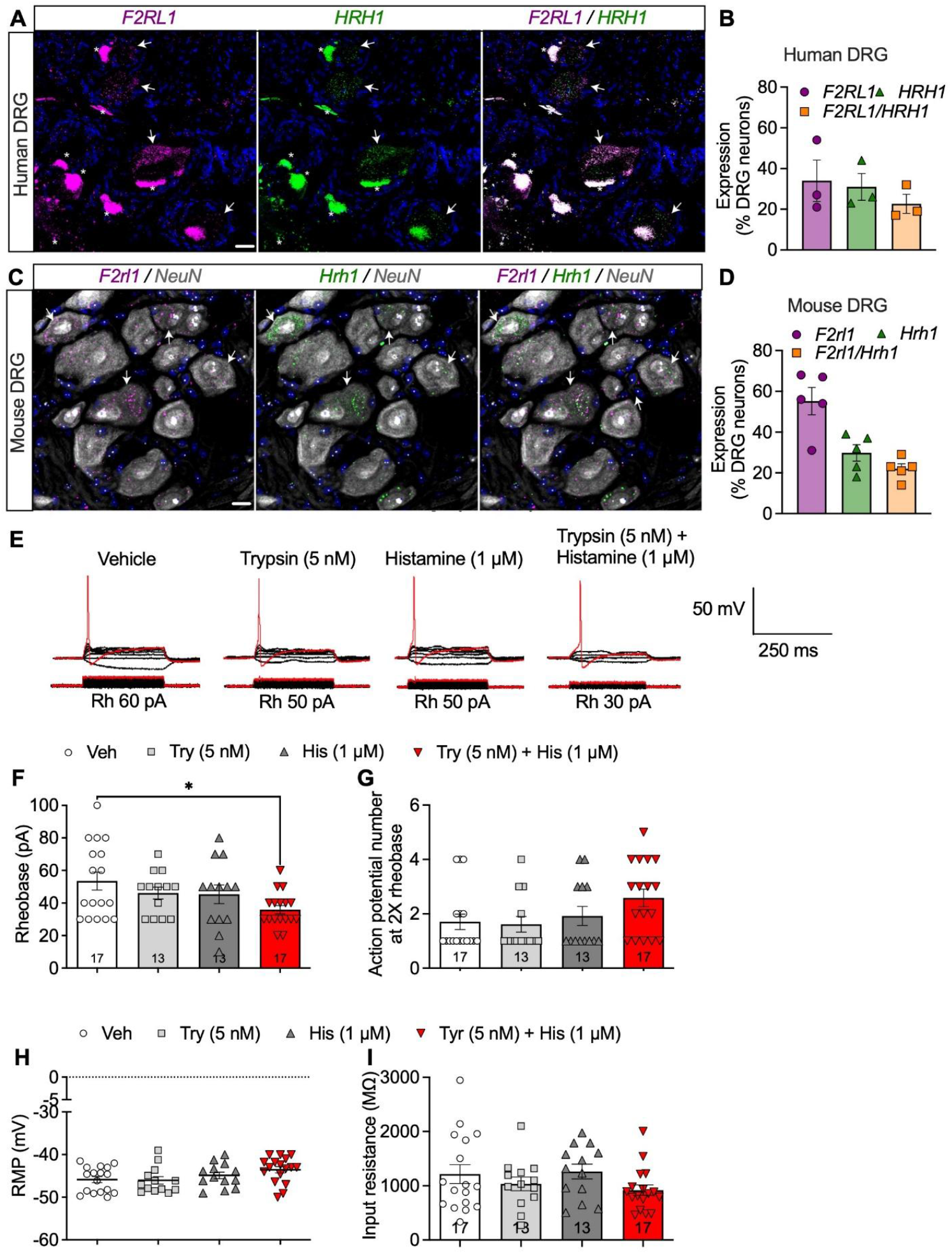
Coexpression and functional synergy of PAR_2_ and H_1_R in DRG neurons. **A-D.** RNAscope *in situ* hybridization localization of mRNA encoding PAR_2_ (*F2RL1, F2rl1*) and H_1_R (HRH1, *Hrh1*) in human (**A, B**) and mouse (**C, D**) DRG neurons, identified in mice by NeuN immunoreactivity. Arrows indicate colocalization of receptors. Asterisks denote lipofuscin fluorescence in human DRG neurons. Representative images from n=3 (human) and n=5 (mice) per group. Scale bars, A 50 µm, C, 20 µm. **E-H**. Electrophysiological recordings of mouse DRG neurons using perforated patch clamp. Representative traces (**E**) and pooled measurements of rheobase (Rh, **F**), action potential number at twice rheobase (**G**), resting membrane potential (RMP, **H**) and input resistance (**I**) of nociceptors treated trypsin (5 nM), histamine (1 µM), both trypsin (5 nM) and histamine (1 µM) or vehicle (Veh). Mean±SEM, data points and numbers in bars indicate the number of neurons from N=6 or 7 mice. \**P*< 0.05, 1-way ANOVA with Tukey’s *post hoc* test.

### PAR_2_ and H_1_R are coexpressed by dorsal root ganglion (DRG) neurons

PAR_2_ and H_1_R are expressed by dorsal root ganglia (DRG) neurons [6, 17, 18]. We used RNAscope *in situ* hybridization to determine if these receptors are expressed in the same neurons in humans and mice. PAR_2_ and H_1_R was mRNA expression was quantified in 3 images per human from 3 donors (N=3; 9 images in total) and 2 images per mouse from 5 subjects (N=5; 10 images in total) (**Table S4**). In humans, *F2RL1* mRNA (PAR_2_) was expressed in 34% of neurons, *HRH1* mRNA (H_1_R) was expressed in 31% of neurons, and *F2RL1* and *HRH1* mRNAs were co-expressed in 23% of neurons; ∼70% of neurons expressing *F2RL1* also expressed *HRH1*) (**Fig. 3A, B, Fig. S2A, C**). In mice, 55% of NeuN-positive neurons expressed *F2rl1*, 30% expressed *Hrh1*, and 22% coexpressed *F2rl1* and *Hrh1*; ∼45% of neurons expressing *F2rl1* also expressed *Hrh1* (**Fig. 3C, D, Fig. S2B, C**). Thus, PAR_2_ and H_1_R are coexpressed in DRG neurons, where they may function synergistically.

### Trypsin and histamine synergistically increase the excitability of DRG nociceptors

To examine potential synergy between PAR_2_ and H_1_R, we measured the effects of trypsin and histamine on the activity of mouse DRG neurons in short-term culture. Neurons were incubated with trypsin or histamine, separately or together, for 15 min, and then washed. As an indicator of excitability, rheobase (minimum input current to elicit action potential) was measured immediately after washing using the perforated patch clamp technique. We previously determined the subthreshold trypsin concentration (5 nM) [17]. The subthreshold histamine concentration (1 µM) was determined by incubating DRG neurons with graded concentrations of histamine (1, 10, 30, 100 µM) for 15 min and measuring rheobase (**Fig. S3**).

Co-application of subthreshold trypsin (5 nM) plus histamine (1 µM) significantly decreased the rheobase (35±2.9 pA) by 34.6±5.5% (F (3, 56)=2.85, [treatment effect],1-way ANOVA, Tukey’s test, *P*<0.05) compared to vehicle (53.5±5.5 pA), consistent with increased excitability (**Fig. 3E, F**). In contrast, trypsin or histamine alone did not alter the rheobase (trypsin: 46.1±3.7 pA; histamine: 45.4±5.7 pA) (**Fig. 3E, F**). Trypsin or histamine did not affect the number of action potentials at twice rheobase, the resting membrane potential or the input resistance, indicating increased excitability occurs without altering baseline electrical properties (**Fig. 3G-I**). Thus, trypsin and histamine act synergistically to sensitize DRG neurons.

### Subthreshold trypsin potentiates histamine-induced sensitization of DRG nociceptors, but not *vice versa*

To determine the relative contributions of PAR_2_ and HRs to the synergistic actions of trypsin and histamine, we treated DRG neurons sequentially with agonists and then measured excitability (**Fig. 4A**). Mouse DRG neurons were preincubated with subthreshold trypsin (5 nM) or histamine (1 µM) for 15 min, washed, and then incubated with the other agonist for 15 min. As a positive control, neurons were co-incubated with both trypsin and histamine for 15 min. As a negative control, neurons were incubated with vehicle. After treatment, neurons were washed and the rheobase and frequency of action potential firing during a depolarization ramp (0-250 pA, 1s) were immediately measured. Simultaneous incubation with trypsin and histamine significantly decreased the rheobase and increased the frequency of action potential firing compared to vehicle, consistent with synergistic signaling and hyperexcitability (**Fig. 4B-D**). In neurons that were pre-incubated with trypsin, histamine significantly decreased rheobase and increased action potential firing compared to vehicle, indicating increased neuronal excitability. In contrast, in neurons that were pre-incubated with histamine, trypsin did not affect rheobase or action potential firing. Thus, PAR_2_ preactivation sensitizes histamine-stimulated excitability whereas HR preactivation does not sensitize trypsin-stimulated excitability. PAR_2_ makes a dominant contribution to trypsin and histamine synergy.

**Figure 4.**
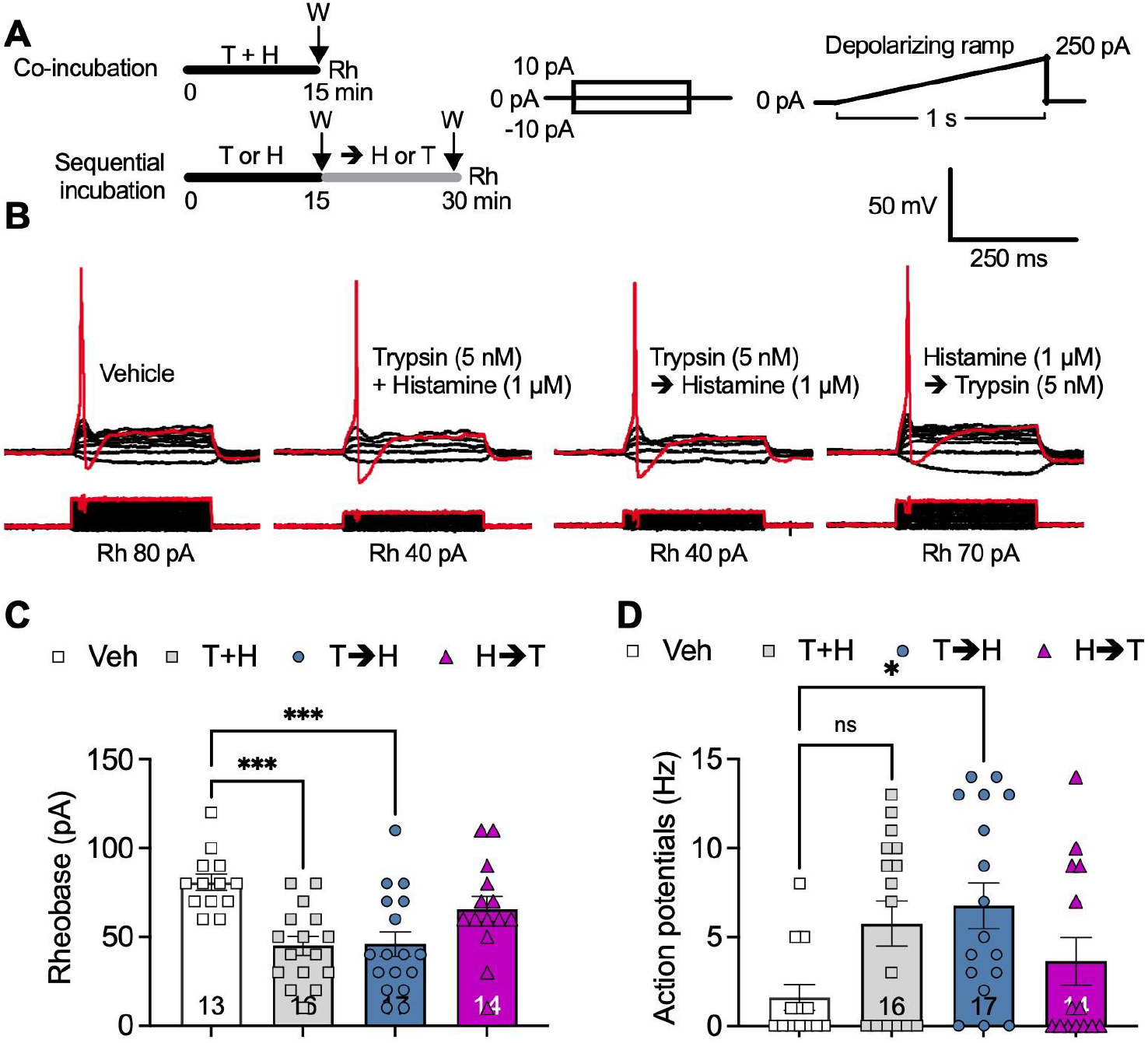
Trypsin-induced potentiation of histamine-stimulated sensitization of DRG neurons. **A.** Experimental protocol. Mouse DRG neurons were incubated with trypsin (T) or histamine (H) individually for 15 min, washed (W) and then incubated with the other agonist for 15 min. Other neurons were co-incubated with trypsin plus histamine (H) (positive control) or were incubated with vehicle (negative control) for 15 min. Rheobase (Rh) was measured during a stepwise current injection protocol (steps: 10 pA, 250 ms, starting at −10 pA) and action potential firing was measured using a ramp protocol. **B-D**. Representative recordings (**B**) and pooled measurements of rheobase (**C**) and action potential firing frequency (**D**) after a depolarizing ramp. Mean±SEM, data points and numbers in bars indicate the number of neurons from N=6-8 mice.\**P*< 0.05, \*\**P*< 0.01, 1-way ANOVA with Tukey’s *post hoc* test (C) and Kruskal-Wallis with Dun’s *post-hoc* test (D).

### PAR_2_ endosomal signaling mediates sensitization of histamine-stimulated nociceptor excitability

Activated PAR_2_ undergoes clathrin-dependent endocytosis and continues to signal from endosomes [16, 17, 18, 19]. Endosomal signaling of PAR_2_ mediates the sustained stimulatory actions of proteases, including those released from mucosal biopsies from IBS-D patients, on nociceptor excitability [17]. We therefore determined whether endosomal signaling of PAR_2_ contributes to the synergistic actions of trypsin and histamine on nociceptor excitability.

To assess the contribution of endocytosis to the excitability of nociceptors, we pre-incubated mouse DRG neurons with the clathrin inhibitor pitstop2 (PS2, 15 µM) or vehicle (control). Neurons were then incubated with subthreshold trypsin (5 nM) plus histamine (1 µM) for 15 min and washed. Rheobase was measured immediately after washing (T=0 min, **Fig. 5A**) or after 15 min recovery (T=15 min, **Fig. 5B**). Trypsin plus histamine decreased rheobase at both T=0 and T=15 min, consistent with an immediate and sustained increase in excitability. PS2 did not affect the immediate increase in excitability (**Fig. 5A**) but reversed the sustained increase in excitability (rheobase: PS2, 69.2±7.4 pA; PS2 vehicle, 40±4.3 pA; (t=3.41, df=19.53), Welch’s t-test, *P*<0.01) (**Fig. 5B**). PS2 also prevented the sustained increase in neuronal excitability induced by the sequential incubation of trypsin and then histamine (rheobase: PS2, 75±7.2 pA; PS2 vehicle, 44±5.8 pA; (t=3.3, df=21.97) Welch’s t-test, *P*<0.01) (**Fig. 5 C**). The dynamin inhibitor dyngo4a (Dy4, 15 µM) also prevented the sustained increase in excitability observed in neurons sequentially incubated with trypsin and then histamine (rheobase: Dy4, 71.8±8.1 pA; Dy4 vehicle, 33±3.9 pA; (t=4.25, df=14.62), *P*<0.001) (**Fig. 5C**). These results suggest that persistent signaling of PAR_2_ in endosomes underlies trypsin-induced sensitization of the effects of histamine on excitability of nociceptors.

**Figure 5.**
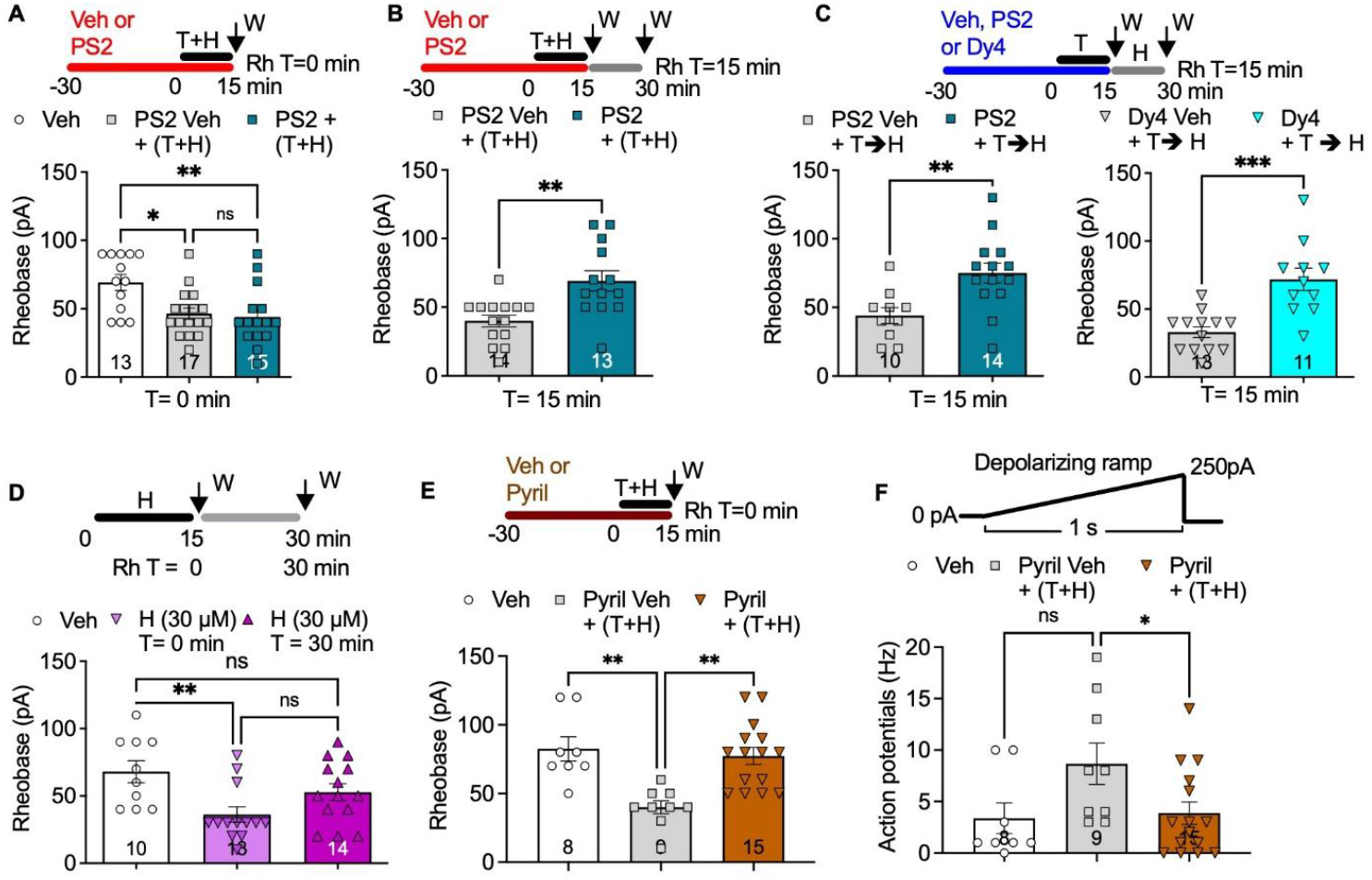
The contribution of endocytosis and the H_1_R to trypsin- and histamine-induced sensitization of DRG neurons. Electrophysiological recordings of mouse DRG neurons. **A, B**. Neurons were preincubated with PS2 (15 µM) or vehicle (Veh, 0.15% DMSO) for 30 min. Neurons were then incubated with trypsin (T, 5 nM) and histamine (H, 1 µM) for 15 min and washed (W). Rheobase (Rh) was measured immediately after washing (T=0 min, A) or after 15 min recovery (T=15 min, B). **C**. Neurons were preincubated with Dy4 (15 µM) or vehicle for 30 min. Neurons were then incubated with trypsin (5 nM) for 15 min, washed and incubated with histamine (1 µM) for 15 min. Rheobase was measured immediately after washing (T=15 min post-trypsin). **D**. Neurons were preincubated to histamine (1 µM) for 15 min and washed. Rheobase was measured **i**mmediately after washing (T=0 min) or after 15 min recovery (T=15 min). **E, F**. Neurons were preincubated with pyrilamine (Pyril, H_1_R antagonist, 1 µM) for 30 min and then incubated with trypsin (5 nM) and histamine (1 µM) for 15 min. Rheobase (E) and action potential firing after a depolarizing ramp (F) were measured at T= 0 min. Mean±SEM, data points and numbers in bars indicate the number of neurons from N=6-9 mice (A), 6-8 mice (B), 6-9 mice (C), 6 mice (D) and 4-7 mice E, F). \**P*< 0.05, \*\**P*< 0.01, \*\**P*< 0.001, 1-way ANOVA with Tukey’s *post hoc* test (A, D and E), Welch’s t test (B, C), and 1-way ANOVA with Dunnett’s *post hoc* test (F).

### Histamine activates the H_1_R to induce immediate but not sustained sensitization of DRG nociceptors

To determine whether histamine causes both an immediate and a sustained sensitization of nociceptors, DRG neurons were incubated with histamine (30 µM) for 15 min and then washed. Rheobase was measured immediately after washout (T=0 min) and after 30 min recovery (T=30 min). At T=0 min, histamine significantly reduced the rheobase compared to control (histamine, 36.1±5.7 pA; control, 68±8.1 pA; F (2, 34)=5.43 [treatment effect], *P*<0.01, one-way ANOVA with Tukey’s test, *P*<0.01) (**Fig. 5D**). In contrast, the rheobase returned to control at T=30 min (52.8±6.2 pA). Thus, although histamine immediately increases the excitability of DRG neurons, the effect is short lived and absent after 30 min. In contrast, trypsin and PAR_2_-induced sensitization is maintained for 30 min.

To identify the HR subtype that mediates synergy with PAR_2_, we incubated DRG neurons with pyrilamine, an H_1_R antagonist, before incubation with subthreshold concentrations of trypsin (5 nM) and histamine (1 µM). Pyrilamine prevented the effects of trypsin and histamine on excitability and action potential firing (**Fig. 5E, F**). Thus, H_1_R activation is required for the synergistic effects of histamine and trypsin on nociceptor excitability.

### Trypsin sensitizes histamine-stimulated activation of H_1_R, β-arrestins and Gαq

To explore the molecular mechanisms underlying trypsin and histamine synergy, we coexpressed PAR_2_ and H_1_R in HEK293 cells, which are commonly used to study GPCR signaling. We examined receptor activation and G protein signaling using enhanced bystander bioluminescence resonance energy transfer (ebBRET) biosensors [23]. ebBRET exploits the endogenous affinity of proteins derived from *Renilla reniformis* to boost the sensitivity (**Fig. 6A**). To measure receptor and β-arrestin activity, we expressed mini (m) mGαq or β-arrestin2 tagged with *Renilla* luciferase (Rluc). mGα are truncated G proteins that diffuse throughout cells to interact with activated conformations of GPCRs [24]. β-arrestins associate with activated GPCRs, including PAR_2_, to mediate receptor desensitization, endocytosis and signaling [15, 16]. To measure Gαq activity, we used a G protein effector membrane translocation assay (GEMTA), which measures the recruitment of Rluc tagged p63-RhoGEF, a Gαq effector [25]. Rluc8-mGαq, β-arrestin2-RlucII or p63-RhoGEF-RlucII were expressed with *Renilla* green fluorescent protein (rGFP) anchored at the plasma membrane (rGFP-CAAX) or early endosomes (tdrGFP-Rab5a). These biosensors were used to monitor activity of PAR_2_, H_1_R, β-arrestin2 and Gαq at the plasma membrane and in early endosomes.

**Figure 6.**
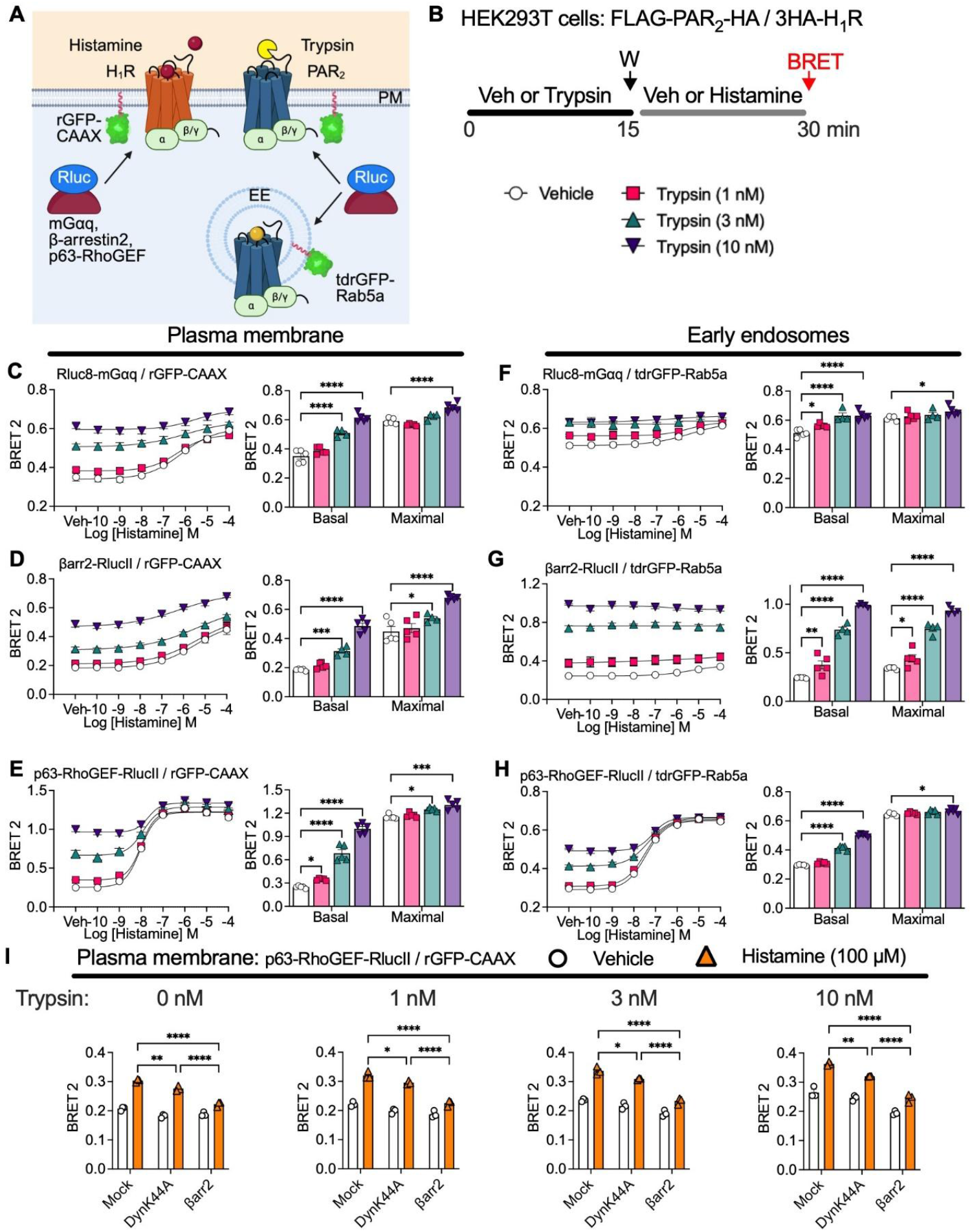
Trypsin and PAR_2_-induced potentiation of histamine-stimulated activation of H_1_R,β-arrestin2 and Gαq. **A.** ebBRET assays of PAR_2_ and H_1_R signaling at the plasma membrane(PM) and in early endosomes (EE). **B**. Experimental protocol. HEK293 cells expressing FLAG-PAR_2_-HA, 3HA-H_1_R and the indicated ebBRET biosensors were preincubated with vehicle (Veh) or trypsin (1, 3, 10 nM) for 15 min, washed (W) and incubated with histamine (0.1 nM-100 µM) for 15 min before ebBRET measurements. **C-E**. Recruitment of Rluc8-mGαq (C), β-arrestin2 (βarr2)-RlucII (D) and p63-RhoGEF-RlucII (E) to the plasma membrane marker rGFP-CAAX. **F-H**. Recruitment of Rluc8-mGαq (F), βarr2-RlucII (G) and p63-RhoGEF-RlucII (H) to the early endosome marker tdrGFP-Rab5a. **I**. Effects of expression of DynK44A or β-arrestin2 on trypsin - induced sensitization of histamine-stimulated recruitment of p63-RhoGEF-RlucII to the plasma membrane marker rGFP-CAAX. Mean±SEM, triplicate measurements, n=3-5 independent experiments. \**P*<0.05, \*\**P*<0.01, \*\*\**P*<0.001, \*\*\*\**P*<0.0001, 2-way ANOVA, Dunnett’s or Šídák’s *post hoc* test.

To replicate the electrophysiological assays in DRG neurons, cells were preincubated with vehicle or trypsin (1, 3, 10 nM) for 15 min, washed and then incubated with graded concentrations of histamine for 15 min, when ebBRET was measured (**Fig. 6B**). In cells preincubated with vehicle, histamine stimulated a concentration-dependent increase in ebBRET between Rluc8-mGαq, β-arrestin2-RlucII or p63-RhoGEF-RlucII and rGFP-CAAX (**Fig. 6C-E**). In vehicle-treated cells, histamine stimulated a concentration-dependent increase in ebBRET between p63-RhoGEF-RlucII and tdrGFP-Rab5a but did not strongly stimulate ebBRET between Rluc8-mGαq or β-arrestin2-RlucII and tdrGFP-Rab5a (**Fig. 6F-H**). These results provide evidence that histamine activates H_1_R and β-arrestin2 at the plasma membrane but not in endosomes, and that histamine activates Gαq at both the plasma membrane and in endosomes. Preincubation with trypsin strongly amplified the basal ebBRET signals and the responses to low histamine concentrations for mGαq, β-arrestin2 and p63-RhoGEF at the plasma membrane (**Fig. 6C-E**) and in early endosomes (**Fig. 6F-H**). In cells preincubated with trypsin, the histamine-induced maximal ebBRET signal for mGαq, β-arrestin2 and p63-RhoGEF was amplified at the plasma membrane and the signal for p63-RhoGEF was also amplified in early endosomes. Thus, trypsin activation of PAR_2_ enhances histamine-stimulated activity of H_1_R, β-arrestin2 and Gαq activity.

To determine whether endosomal signaling of PAR_2_ contributes to H_1_R sensitization, we overexpressed dominant negative dynamin K44A (DynK44A), which inhibits PAR_2_ endocytosis and endosomal signaling [18]. We measured histamine (100 µM)-stimulated p63-Rho-GEF ebBRET signals at the plasma membrane in cells pretreated with vehicle or trypsin (1, 3, 10 nM). DynK44A inhibited the ability of trypsin to amplify histamine-stimulated p63-Rho-GEF ebBRET signals when compared to mock transfected control cells (**Fig. 6I**). These results align with the inhibitory effects of clathrin and dynamin inhibitors on trypsin-induced sensitization of histamine-stimulated excitation of DRG neurons (**Fig. 5**).

PAR_2_ is a class B GPCR that exhibits sustained high affinity interactions with β-arrestins in endosomes [16, 26, 27]. H_1_R is a class A GPCR that exhibits low affinity and transient interactions with β-arrestins [28]. Certain class B GPCRs sequester β-arrestins in endosomes and thereby impede β-arrestin-mediated desensitization and endocytosis of class A GPCRs [29]. In support of these reports, overexpression of β-arrestin2 inhibited the ability of trypsin to amplify histamine-stimulated p63-Rho-GEF ebBRET signals when compared to mock transfected control cells (**Fig. 6I**). These results suggest that PAR_2_ endocytosis and endosomal signaling, coupled with the sequestration of β-arrestin in endosomes with PAR_2_, amplify H_1_R signaling and mediate the synergistic effects of trypsin and histamine.

### Histamine does not sensitize trypsin-stimulated activation of PAR_2_, β-arrestins and Gαq

To determine whether histamine activation of the H_1_R could similarly potentiate trypsin-induced PAR_2_ activation, we pre-incubated HEK293 cells expressing PAR_2_ and H_1_R with vehicle or histamine (0.1, 1, 10 µM) for 15 min, washed, stimulated cells with graded concentrations of trypsin for 15 min, and then measured ebBRET (**Fig. 7A**). In vehicle-treated cells, trypsin stimulated a concentration-dependent increase in ebBRET between Rluc8-mGαq, β-arrestin2-RlucII or p63-RhoGEF-RlucII and rGFP-CAAX (**Fig. 7B-D**) and tdrGFP-Rab5a (**Fig. 7E-G**). These results show that trypsin stimulates mGαq and β-arrestin2 recruitment to the PAR_2_ at the plasma membrane and in early endosomes and are consistent with mGαq activation at the plasma membrane and in early endosomes. Histamine pre-incubation increased the basal signal of mGαq and p63-RhoGEF at the plasma membrane ((**Fig. 7B, E**) and in early endosomes (**Fig. 7D, G**) compared to vehicle. The highest concentration of histamine (10 µM) slightly increased the maximal trypsin-stimulated signal for p63-RhoGEF at the plasma membrane and in early endosomes. However, histamine had no effect on the maximal trypsin-stimulated signals for mGαq or β-arrestin2 at the plasma membrane and in early endosomes. Thus, in contrast to trypsin, which markedly enhances basal and histamine-stimulated H_1_R, mGαq and β-arrestin2 activities, histamine pretreatment had little or no effect on trypsin-stimulated activity of PAR_2_, β-arrestin2 or Gαq. These findings likely explain why trypsin pretreatment sensitizes histamine responses, but not *vice versa*.

**Figure 7.**
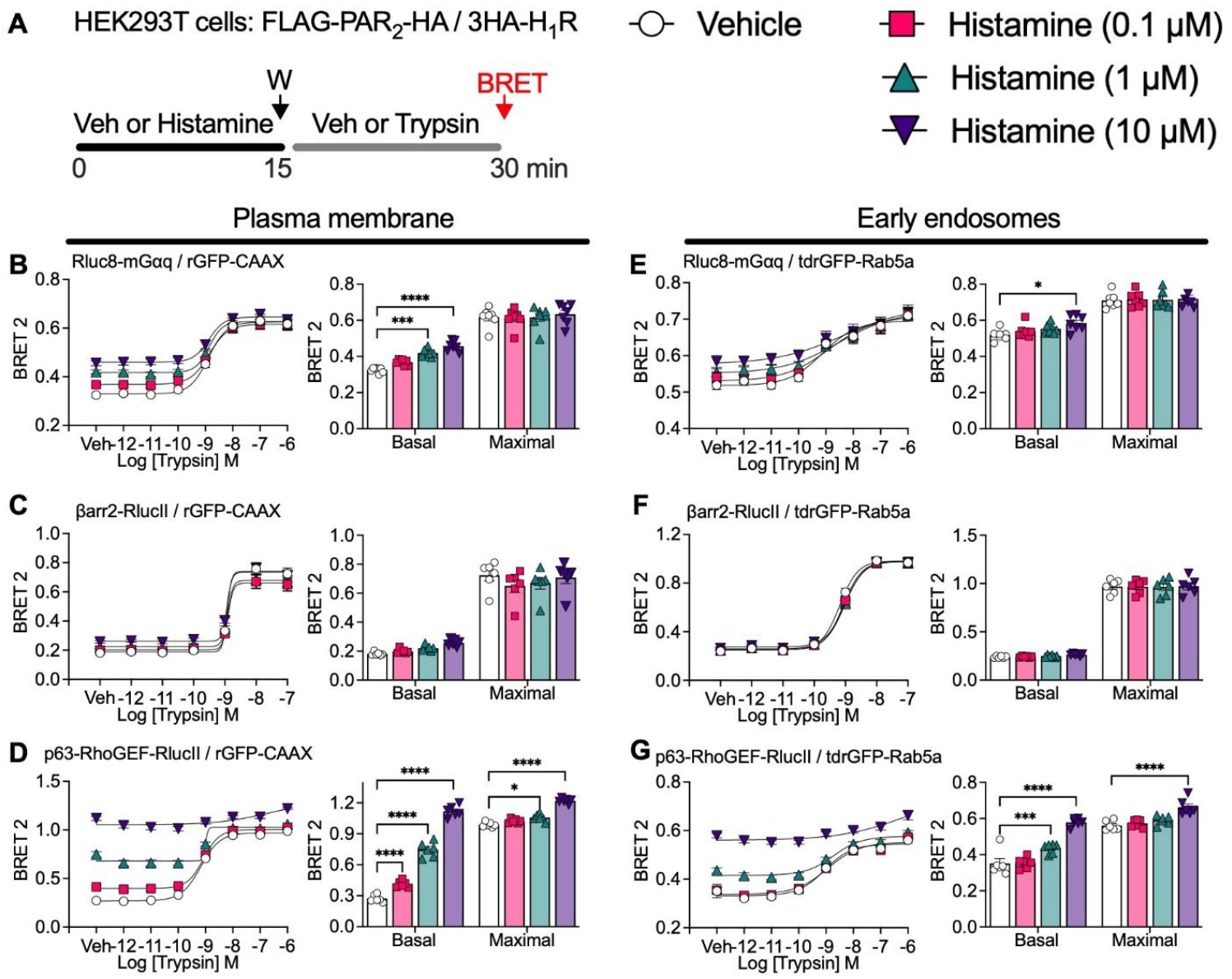
Inability of histamine and H_1_R to potentiate trypsin-stimulated activation of PAR_2_,β-arrestin2 and Gαq. **A.** Experimental protocol. HEK293 cells expressing FLAG-PAR_2_-HA, 3HA-H_1_R and the indicated ebBRET biosensors were preincubated with vehicle (Veh) or histamine (0.1, 1, 10 µM) for 15 min, washed (W) and incubated with trypsin (1 pM-1 µM) for 15 min before ebBRET measurements. **B-D**. Recruitment of Rluc8-mGαq (B), β-arrestin2 (βarr2)-RlucII (C) and p63-RhoGEF-RlucII (D) to the plasma membrane marker rGFP-CAAX. **E-G**. Recruitment of Rluc8-mGαq (E), βarr2-RlucII (F) and p63-RhoGEF-RlucII (G) to the early endosome marker tdrGFP-Rab5a. Mean±SEM, triplicate measurements, n=6-7 independent experiments. \**P*<0.05,\*\**P*<0.01, \*\*\**P*<0.001, \*\*\*\**P*<0.0001, 2-way ANOVA, Dunnett’s *post hoc* test.

## Discussion

Our results provide evidence that trypsin and histamine, which are upregulated in the colon of IBS-D patients, act in concert to enhance colonic nociceptor mechanosensitivity and amplify and sustain DRG nociceptor excitability. PAR_2_ and H_1_R respectively mediates the synergistic effects of trypsin and histamine. PAR_2_ is the dominant partner because preactivation of PAR_2_ sensitizes the H_1_R but not *vice versa*. The irreversible mechanism of PAR_2_ activation and persistent endosomal signaling underly the sustained synergistic effects of trypsin because endocytosis inhibitors prevent synergism. Measurements of PAR_2_, H_1_R, β-arrestin2 and Gαq signaling in HEK293 cells indicate that PAR_2_ endosomal signaling potentiates histamine-stimulated activation of the H_1_R and its β-arrestin2 and Gαq effectors. Our results provide evidence that antagonism of PAR_2_ will blunt the pronociceptive actions not only of proteases but also of histamine and possibly other pronociceptive factors that signal pain in a cooperative manner with PAR_2_.

### PAR_2_ and H_1_R cooperate to sensitize nociceptors

Intracolonic administration of IBS-DS FS, which contains proteases and histamine [12], increased basal activity and mechanosensitivity of colonic afferent nerves of mice. Mechanosensitivity was enhanced at higher intracolonic pressures, consistent with the sensitization of high-threshold nociceptors. Our findings align with reports of sensitization of mouse colonic afferents nerves by IBS-derived neuroactive factors [5, 7, 10, 11, 12]. PAR_2_ and H_1-4_R antagonists, independently or together, prevented IBS-D FS-induced mechanosensitivity. Coadministration of subthreshold trypsin and histamine replicated the effects of IBS-D FS. These combined results support the hypothesis that proteases and histamine, which are produced by immune cells [3, 4] and colonic bacteria [11, 12, 13, 30], act synergistically to potentiate excitatory signaling of nociceptors in IBS-D, contributing to the abdominal pain in patients. Although we cannot exclude the involvement of other mediators, our findings highlight a major synergy between PAR_2_ and H_1-4_R in signaling pain.

Our results show that PAR_2_ and H_1_R mRNAs are co-expressed in ∼20% of human and mouse DRG neurons. Thus, proteases and histamine could act simultaneously on the same neuron to enhance pain signaling. Although we did not specifically assess PAR_2_ and H_1_R expression in DRG neurons innervating the colon, recordings from colonic afferent nerves provide evidence for the presence of these receptors in colonic nociceptors. Patch clamp recordings of DRG provide functional evidence of receptor coexpression. Trypsin and histamine activated PAR_2_ and H_1_R, respectively, to sensitize neurons with properties of nociceptors (*i*.*e*., <30 pF capacitance, small diameter), consistent with previous studies of isolated DRG neurons and *ex-vivo* recordings from mouse colon [6, 17, 22, 31, 32]. Notably, the combined application of subthreshold trypsin and histamine increased the excitability of DRG nociceptors, which reinforces the proposal that PAR_2_ and H_1_R are coexpressed and function synergistically to sensitize nociceptors.

### Mechanisms of PAR_2_ and H_1_R cooperative signaling

The convergence of common pathways of PAR_2_ and H_1_R signaling likely underlies the synergistic actions of trypsin and histamine on nociception (**Fig. 8**). Since both PAR_2_ and H_1_R couple to Gαq, leading to activation of protein kinase C and sensitization of voltage-gated and transient receptor potential ion channels [6, 33, 34], convergence could occur at multiple levels.

**Figure 8.**
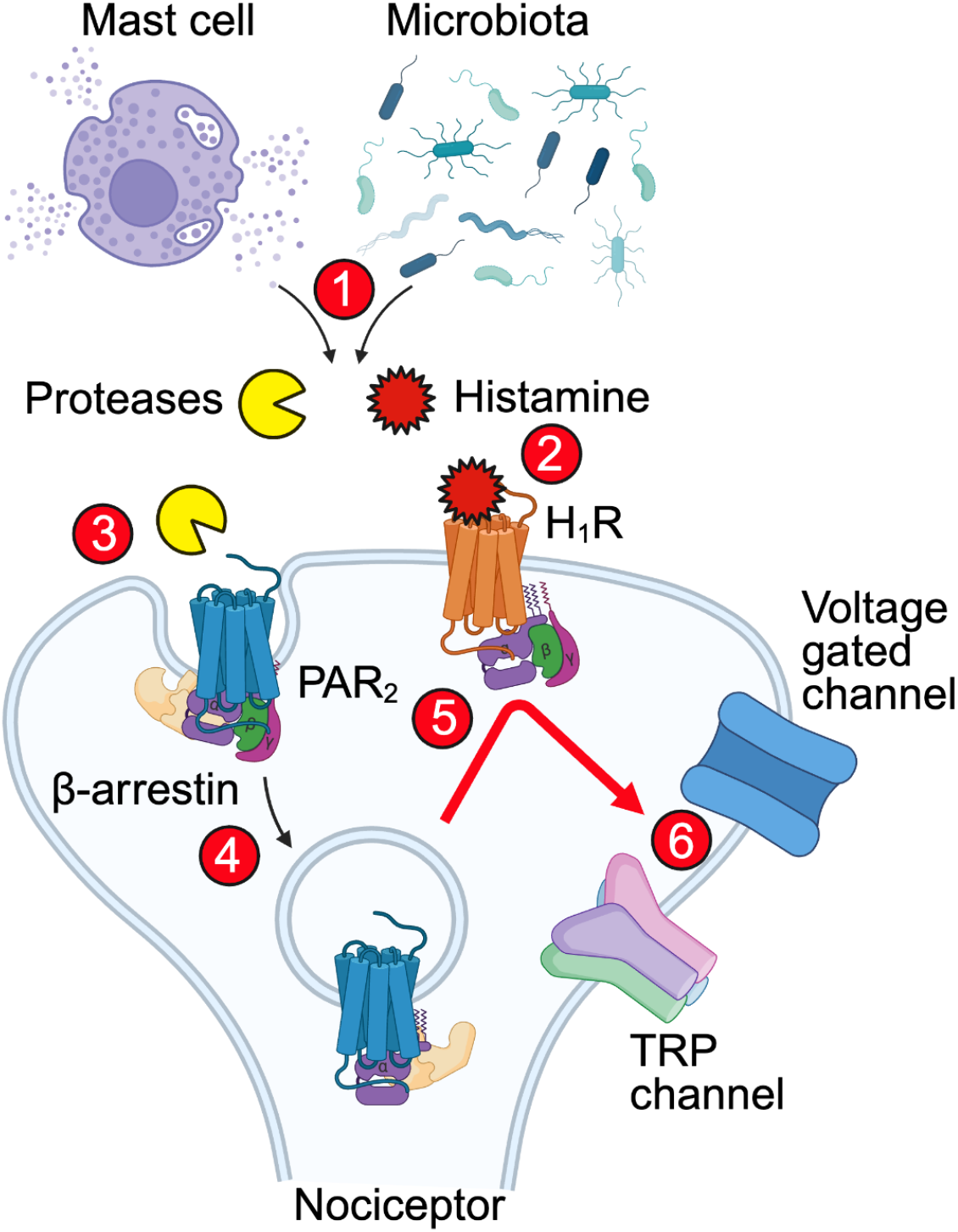
Hypothesized mechanism by which endosomal signaling of PAR_2_ amplifies and sustains H_1_R signaling at the plasma membrane to maintain the hyperresponsiveness of colonic nociceptors and cause recurrent IBS pain. 1. Mast cells and bacteria in the colon of IBS patients secrete histamine and proteases. Histamine reversibly binds to and activates the H_1_R, and proteases irreversibly cleave and activate PAR_2_ at the plasma membrane of nociceptors innervating the colon. **2**. H_1_R couples to Gαq at the plasma membrane and transiently interacts with β-arrestins, which desensitize and thereby terminate H_1_R signaling. **3**. PAR_2_ couples to Gαq and β-arrestins at the plasma membrane. **4**. PAR_2_ undergoes clathrin-mediated endocytosis and exhibits sustained interaction with Gαq and β-arrestins in endosomes. **5**. By sequestering β-arrestins, PAR_2_ amplifies and sustains H_1_R signaling. **6**. Amplified and sustained H_1_R signaling from the plasma membrane and prolonged PAR_2_ signaling from endosomes together sensitize voltage-gated and transient receptor potential (TRP) ion channels and thereby magnify and maintain the sensitization of nociceptors to cause recurrent IBS pain.

To determine the relative contributions of PAR_2_ and H_1_R signaling to the synergistic actions of trypsin and histamine, we challenged nociceptors sequentially with trypsin or histamine and then examined excitability. Whereas preactivation of PAR_2_ enhanced H_1_R-induced excitability, pretreatment with histamine did not magnify responses to trypsin. Measurements of the activity of PAR_2_, H_1_R and Gαq and β-arrestin2 effectors in subcellular compartments of HEK293 cells reinforced these findings. Preactivation of PAR_2_ enhanced histamine-stimulated activity of H_1_R and β-arrestin2 at the plasma membrane and Gαq at the plasma membrane and in endosomes (**Fig. 6**). Conversely, the preactivation of H_1_R had no effect on trypsin-stimulated activation of PAR_2_ or β-arrestin2, with only modest effects on Gαq activity (**Fig. 7**). The dominant role of PAR_2_ accords with the irreversible mechanism of proteolytic activation and its known ability of PAR_2_ to continue to signal from endosomes [16, 17, 18, 19], which was confirmed by the present results showing trypsin-stimulated activity of PAR_2_, Gαq and β-arrestin2 in early endosomes. We found that trypsin caused an immediate and a sustained sensitization of nociceptors, in agreement with previous reports [17]. The observations that dominant negative DynK44A, which inhibits endocytosis and endosomal signaling of PAR_2_ [17], and endocytosis inhibitors suppressed trypsin-induced sensitization of the H_1_R fsupports involvement of sustained endosomal signaling of PAR_2_. In contrast, although histamine had an immediate sensitizing effect, the response was short-lived and was not maintained after agonist removal.

Differences in the activation, signaling, trafficking and regulation of PAR_2_ and H_1_R may account for their different relative contributions to the synergistic effects of trypsin and histamine in nociceptors. Proteases cleave and irreversibly activate PAR_2_, a class B GPCR that exhibits sustained high affinity interactions with β-arrestins, internalizes and continues to signal from endosomes, leading to persistent nociception [16, 17, 18, 19, 26, 27]. In contrast, histamine reversibly binds to the H_1_R, a class A GPCR that transiently interacts with β-arrestins, leading to desensitization of signaling without formation of stable complexes with β-arrestins in the endosomes [9, 28, 35]. Histamine failed to strongly stimulate H_1_R and β-arrestin activity in endosomes of HEK293 cells, in line with these reports. Suppression of β-arrestin2 and GPCR kinase prevents H_1_R desensitization, suggesting that β-arrestin2 recruitment and sequestration by other co-expressed GPCRs might enhance H_1_R signaling [35, 36] (**Fig. 8**). In contrast depletion of β-arrestin may prevent PAR_2_ endocytosis and therefore PAR_2_ endosomal signaling [16, 26]. The finding that overexpression of β-arrestin2 abrogated trypsin-induced sensitization of H_1_R activation of Gαq supports this proposal.

In addition to the H_1_R, PAR_2_ can potentiate responses to other GPCRs, receptor tyrosine kinases and ligand-gated ion channels that control sensation [7, 37, 38, 39, 40]. Although the underlying mechanisms for these interactions may differ, our work provides evidence that PAR_2_ can sensitize multiple signaling pathways that lead to pain.

### Limitations

We cannot exclude the possibility that proteases and histamine act on cells other than nociceptors in the colon to induce pain. PAR_2_ and H_1_R are expressed by epithelial and immune cells that have been implicated in IBS pain [14, 41]. PAR_2_ activation in colonic epithelial cells disrupts the barrier leading to nociception in mice [18]. Histamine induces mast cell accumulation and degranulation, causing visceral hypersensitivity [11, 42]. We did not characterize co-expression of *F2rl1* and *H1R* in colonic nociceptors and our functional studies in the DRG are not selective for colonic neurons. Although analysis of signaling in HEK293 cells provides insights into mechanisms of synergism, analysis of signaling in colonic nociceptors will be required to definitively delineate the mechanisms of cooperative signaling in functionally relevant cells. We used genetic and pharmacological approaches to determine the contribution of endocytosis to synergistic PAR_2_ and H_1_R interaction. However, Dy4 and PS2 have non-specific effects on endocytosis [43]. The use of nanoparticle-encapsulated PAR_2_ antagonists, which selectively inhibit endosomal signaling [19], will be required to more definitively determine the role of endosomal signaling in IBS pain.

### Therapeutic implications

Our findings illustrate the multimodal nature of pain, whereby seemingly redundant receptors and signaling pathways converge to amplify and sustain chronic pain. They have implications for the identification of predictive biomarkers and the development of therapies. Although proteolytic activity and histamine levels are elevated in mucosal biopsies and fecal samples from IBS-D patients, whether the concentrations are sufficient to excite nociceptor terminals in the colon is unknown. Analysis of colonic biopsies and FS from IBS-D patients indicates that tryptase is present at picomolar concentrations and histamine is in the nanomolar range [3, 12], which are lower than those used to activate PAR_2_ and H_1_R in the current study, yet still sufficient to sensitize nociceptors [4, 5, 6, 11, 12, 17]. Proteases, histamine and other neuroactive factors from immune cells and the microbiome likely act synergistically to induce and sustain pain even at concentrations that, individually, are inactive. Thus, profiling a panel of biomarkers rather than measuring a single mediator would be required to understand mechanisms and identify therapeutic targets. The multimodal characteristics of pain, in which a single nociceptor can express multiple receptors that induce hyperexcitability, poses a challenge for drug development. However, the identification of dominant receptors and points of convergence of signaling pathways that underlie synergistic mechanisms can inform the development of effective therapies. Our results provide evidence that antagonists of PAR_2_ can block the pronociceptive actions of proteases, histamine and possibly other neuroactive factors in IBS-D patients. An in-depth analysis of pain mediators that are upregulated in painful diseases together with comprehensive analysis of the expression, activity and cooperative signaling of receptors on nociceptors will be required to identify more effective treatments for chronic pain.

## Methods

See supplement.

### Patients

Queen’s University Health Sciences and Affiliated Teaching Hospitals Research Ethics Board and the Institutional Review Board of the University of Cincinnati approved human studies. Pain severity was assessed by IBS-SSS [44] and PROMIS abdominal pain raw data score [45].

### FS preparation

Fecal samples were diluted and homogenized in Krebs solution, centrifuged and sterile filtered.

### Mice

Queen’s University Animal Care Committee and New York University Institutional Animal Care and Use Committee approved studies on C57BL/6 mice.

### Extracellular colonic afferent recording

The distal colon was prepared for extracellular recording from colonic afferents in response to distention [12].

### DRG neuron patch-clamp recording

DRG neurons were dispersed and rheobase and action potential firing were measured by patch-clamp [17].

### RNAscope

DRG were collection of de-identified organ donors and from mice. mRNA transcripts were localized using the RNAscope.

### ebBRET

HEK293 cells were transfected with ebBRET biosensors to measure recruitment of mGα, β-arrestin2 [23], and p63-RhoGEF [25] to the plasma membrane and early endosomes.

### Statistics

Results were tested for normality using D’Agostino-Pearson omnibus normality test. Results were analyzed by t-test or ANOVA with Šidák’s, Tukey’s, Dunnett’s or Dunn’s *post hoc* test. Non-parametric results were analyzed using Wilcoxon and Kruskal-Wallis tests with Dunn’s *post hoc* test. Results are expressed as mean±SEM with significance *P*<0.05.

## Author contributions

NNJV, SD, DER, AEL, SJV and NWB designed the experiments. NNJV, BS, DAG and MFP completed the experiments and analyzed the results. NNJV, SJV and NWB wrote the manuscript. BLS, DER, AEL, SJV and NWB obtained funding.

## Acknowledgments

Supported by NIH grants R01DE029951(NWB, BLS), RM1DE033491 (NWB, BLS), DoD grant W81XWH2210239 (NWB, BLS) and project grants from the CIHR (DER, AEL, SJV).

