## Supplementary material for "ENDOSOMAL SIGNALING OF PROTEASE-ACTIVATED RECEPTOR-2 AMPLIFIES HISTAMINE-INDUCED PAIN OF IRRITABLE BOWEL SYNDROME": PAR2 H1R Supplement 01.29.2026.pdf

### Methods

**Patients.** The Queen's University Health Sciences and Affiliated Teaching Hospitals Research Ethics Board (protocol number DMED 2047-17) approved studies on IBS patients. Fecal samples were collected from individuals enrolled in the inflammation, microbiome, alimentation, gastrointestinal and neuropsychiatric effects (IMAGINE) study in Canada. Patients that fulfilled the Rome IV criteria for IBS-D were enrolled. After sample donation, patients completed the IBS Severity Score System (IBS-SSS) and the Patient-Reported Outcomes Measurement Information System (PROMIS) abdominal pain score questionnaire to stratify patients into high, moderate or low pain categories. Samples from three female patients with IBS-D with high IBS-SSS and PROMIS classification were used (**Table S1**). Patients were instructed to collect samples in designated containers to avoid contamination by water or urine. The samples were stored at -20 °C and promptly delivered to Kingston General Hospital, where they were stored at -80 °C and used for FS preparation.

**Pain severity assessment.** The IBS-SSS comprises 5 questions each with a scale of 0 (not applicable) to 100 (very applicable) [1] (**Table S2**). The IBS-SSS was scored out of 500 and graded as low (75-174), moderate (175-299) or severe (300-500). The PROMIS abdominal pain score questionnaire comprises 5 questions each with a scale of 0 (no pain) to 5 (maximal pain) [2] (**Table S3**). The PROMIS was scored out of 25 and the raw data scores were graded as low pain (5-12), moderate pain (13-18) or high pain (19-25).

**FS preparation.** Fecal samples were diluted, mixed and homogenized in sterilized Krebs solution (mM: 118.4 NaCl, 24.9 $NaHCO_3$, 1.2 $MgSO_4$, 1.2 $KH_2PO_4$, 11.7 glucose, 1.9 $CaCl_2$, pH 7.4) in a ratio of 0.5 g per 4 mL. Samples were centrifuged (10,000 g, 10 min, 4°C) and the supernatant was filtered through a 40 µm cell strainer and then through a sterile 0.45 µm and 0.2 µm syringe filter. FS was aliquoted and stored at -80 °C.

**Mice.** The Queen's University Animal Care Committee (#2020-2011) and the New York University Institutional Animal Care and Use Committee (PROTO201900106) approved animal studies. Experiments were consistent with the Canadian Council of Animal Care and the -ARRIVE 2.0 guidelines. Male and female C57BL/6 mice (8-12 weeks) were obtained from Queens University and the Jackson laboratory. Mice were maintained in a light-controlled (12-h light/dark cycle) and temperature-controlled (22°C) environment with access to food and water *ad libitum*.

***Ex vivo* extracellular recordings from colonic afferent nerves.** Mice were euthanized by isoflurane inhalation. For extracellular recording, the distal colon, defined as the 3 cm segment immediately proximal to the pelvic brim, was excised along with the associated neurovascular bundle proximal to the inferior mesenteric ganglion and distal aorta [3]. The resected segment was placed in an organ bath and continuously perfused with carbogenated (95% $O_2$ / 5% $CO_2$) Krebs solution, pH 7.4 at 34°C. The colon was cannulated at both ends. The proximal end was connected to an intraluminal infusion pump, and the Krebs solution was perfused through the lumen at 0.2 mL/min. The distal end was connected to an intraluminal pressure transducer (NL108; Digitimer, Welwyn Garden City, UK). The lumbar splanchnic nerve was identified, its distal end was isolated and suctioned into a glass electrode attached to a Neurolog headstage

(NL100, Digimiter). The recorded signals were amplified using an A.C. preconditioned amplifier (NL104) set to 10,000x and filtered (10-5000 Hz, sampling frequency 20 KHz, NL125 band-pass filter). The analog signal was digitized using a Micro 1401 MKII interface and recorded and analyzed on CED Spike2 6.1 software. To minimize smooth muscle activity, the Krebs solution was supplemented with nifedipine (3 µM, L-type calcium channel blocker), atropine (5 µM, muscarinic acetylcholine receptor antagonist) and indomethacin (3 µM, cyclooxygenase inhibitor to prevent the formation of inhibitory prostaglandins).

After setup, the preparation was rested for 15 min. The colon was then distended with a pressure ramp by closing the outflow drain until intraluminal pressure reached 60 mm Hg. The drain was then opened, returning the pressure to baseline, and the preparation was rested for 15 min. The preparation was considered stable when the afferent frequency responses to 3 consecutive distensions were within 20%. Basal firing was determined by averaging action potential firing during the 120 s before the final reproducible distension. The last reproducible distension was considered the control distension.

Once reproducibility was established, the lumen of the colon was perfused with FS (diluted 1:2 in Krebs), histamine (30, 300 µM), trypsin (30, 50 µM) or a combination of subthreshold concentrations of histamine (30 µM) and trypsin (30 µM) for 20 min. Some preparations were pre-treated with antagonists of $PAR_2$ (10 µM GB83) or histamine receptors (1 µM pyrilamine, 10 µM ranitidine, 30 nM clobenpropit, 1 µM JNJ7777120) for 20 min before perfusion with FS and antagonists for an additional 20 min. At the end of the treatment period, the colon was distended in the presence of the treatment. The colon was then perfused with Krebs solution to remove antagonists and FS and recovered for 15 min. Basal firing was measured for 120 s and the colon was distended to measure post-treatment responses.

Data were analyzed offline. All units of a recording were analyzed by extracting all spike templates representing neural units in that recording. The low threshold units were disregarded because they are not suspected to mediate nociception. The units from the different recordings under the same treatment condition were then grouped according to the individual unit classification. Treatment- and washout-frequency distension responses were normalized to percentage of maximum control distension.

**Dissociation of DRG neurons for patch clamp recordings.** DRG (T11 - L3) were excised, digested and cultured overnight [4]. Briefly, DRG neurons were incubated with collagenase IV (1mg/mL, Worthington) and dispase (4 mg/mL, Roche) (10 min, 37°C), washed and dissociated by trituration with a fire-polished Pasteur pipette. Neurons were plated onto laminin- (0.017 mg/mL) and poly-D-Lysine- (2 mg/mL) -coated glass coverslips. Cells were incubated in F12 medium (Sigma, cat N6658) containing 10% of fetal calf serum, penicillin and streptomycin (100 U/mL and 0.1 mg/mL, Sigma cat P4333) in a humidified chamber (95% air, 5% $CO_2$, 16 h, 37 °C).

**Patch clamp recording of nociceptors.** Changes in excitability of small-diameter (<30 pF capacitance) DRG neurons with properties of nociceptors were quantified by measuring rheobase (minimum input current to elicit an action potential firing) and action potential firing at twice rheobase in current clamp mode using a stepwise protocol (steps: 10 pA, 250 ms, starting at -10 pA) and action potential number during a depolarization ramp (0 to 250 pA, 1 second) by whole-cell perforated patch-clamp recordings using Amphotericin B (240 µg/mL) [4]. Resting membrane potential was measured and the input resistance was determined by measuring the voltage deflection in response to a single injection of 10 pA hyperpolarizing current. Only neurons with resting membrane potential more negative than -40 mV were analyzed. Recordings were made using Axopatch 200B amplifiers, digitized by Digidata 1550B and stored and processed using pClamp 11 software (Molecular Devices). The recording chamber was continuously perfused with external solution at 2 mL/min. External solution was (mM): 140 NaCl, 5 KCl, 10 HEPES, 10 D-glucose, 1 $MgCl_2$, 2 $CaCl_2$; pH to 7.4 with 3 M NaOH. Pipette solution was (mM): 110 K-gluconate, 30 KCl, 10 HEPES, 1 $MgCl_2$, 2 $CaCl_2$; pH 7.25 with 1 M KOH.

In some experiments, neurons were incubated for 15 min with vehicle (control), trypsin (5 nM), histamine (1, 10, 30, 100 µM) or both trypsin (5 nM) and histamine (1 µM). Neurons were washed and rheobase was measured immediately (T=0 min) or 30 min later (T=15 min). In separate experiments, neurons were challenged with trypsin (5 nM, 15 min) or histamine (1 µM, 15 min) and then washed. Trypsin-incubated neurons were then challenged with histamine (1 µM, 15 min) and histamine-incubated neurons were challenged with trypsin (5 nM, 15 min), or neurons were challenged with vehicle (control). Rheobase was measured at T=0 min or T=15 min. To determine whether clathrin-mediated endocytosis contributes to sensitization, neurons were preincubated with the clathrin inhibitor pitstop2 (15 µM, 30 min) or the dynamin inhibitor dyngo4a (15 µM, 30 min) to prevent receptor endocytosis or the H1R antagonists, pyrilamine (1 µM, 30 min), before agonists. To evaluate the ability of histamine receptors to evoke sustained hyperexcitability, neurons were challenged with 30 µM histamine for 15 min and then washed, then rheobase was measured at T= 0 or T= 30 min after.

**Collection of human tissue for RNAscope.** DRG collection from de-identified organ donors was reviewed by the Institutional Review Board of the University of Cincinnati (IRB #00003152, Study ID 2015-5302) and deemed exempt. DRG were collected from 3 de-identified organ donors. Donor information is provided in **Table S4**. Lumbar DRG were recovered within 90 min of cross-clamp, transported in N-methyl-D-glucamine (NMDG) medium to the laboratory, and immediately cleaned and dissected to remove connective tissue. DRG were sectioned longitudinally in half, post-fixed in 4% paraformaldehyde in PBS overnight at 4°C, cryoprotected in 30% sucrose (24 h, 4°C), and embedded in optimal cutting temperature (OCT) compound (Tissue-Tek). Frozen sections (18-20 µm) were mounted onto Superfrost Plus slides (Fisher, Suwanee, GA), air-dried (15 min) and stored (-20°C).

**Collection of mouse tissue for RNAscope.** Mice were anesthetized (5% isoflurane) and perfused through the ascending aorta with PBS and then 4% paraformaldehyde in PBS. Lumbar DRG were removed, fixed in 4% paraformaldehyde in PBS (4 h, 4 °C), cryoprotected with 30% sucrose (24 h, 4 °C), and embedded in OCT. Frozen sections (10 µm) were mounted onto Superfrost Plus slides, air-dried (15 min) and stored (-20°C).

**RNAscope *in situ* hybridization and immunofluorescence.** mRNA transcripts were localized in human and mouse DRG using the RNAscope system (Advanced Cell Diagnostics) following the manufacturer's instructions for fresh-frozen tissue, except for the omission of the first on-slide fixation step. Probe hybridization and detection used the Multiplex Fluorescent Kit v2 according to the manufacturer's protocol. Probes specific for human (Hs): *Hs-F2RL1* ($PAR_2$, #319861-C1) and *Hs-HRH1* ($H_1R$, #416501-C3), and mouse (Mm): *Mm-F2rl1* ($PAR_2$, #446931-C3) and *Mm-Hrh1* ($H_1R$, #417541-C1) were used. Human and mouse tissues were incubated with TSA Vivid™ Fluorophore 520 (1:1500, Advanced Cell Diagnostics, cat#323271) and TSA Vivid™ Fluorophore 650 (1:1500, Advanced Cell Diagnostics, cat#323273) for fluorescence detection. For the detection of neurons in mouse tissues, the hybridized slides were blocked and incubated overnight at 4°C with guinea pig anti-NeuN antibody (1:500; EMD Millipore, Cat# N90). Slides were then washed and incubated for 1 h at room temperature with goat anti-guinea pig Alexa Fluor® 647 (1:1000). Slides were washed and counterstained with DAPI (1 µg/mL, 5 min) and mounted with ProLong® Gold Antifade Mountant (Thermo Fisher). Sections were imaged using a Leica SP8 confocal microscope equipped with an HCX PL APO 20× objective (human sections) and HCX PL APO 40x (NA 1.30) oil objective (mouse sections). Sections were observed using a Leica SP8 confocal microscope with HCX PL APO 20x or HCX PL APO 40x (NA 1.30) oil objectives (Wetzlar, Germany). A total of 3 images (20× magnification) were analyzed per human sample (N=3; 9 images in total) and 2 images (40× magnification) per mouse (N=5; 10 images in total). The percentage of hybridized positive neurons was quantified and normalized to the total number of human neurons identified by morphology or NeuN+ neurons for mice.

**ebBRET.** HEK293 cells were cultured in DMEM supplemented with 10% fetal bovine serum (FBS), 1% penicillin and streptomycin (100 U/mL and 0.1 mg/mL, Sigma cat P4333). Cells in suspension were transiently transfected at a density of 0.4 million cells per mL using 25 kDa linear polyethylenimine (Polysciences; 4:1 PEI:DNA) and seeded in 96-well microplates (Greiner; #655083) (100 µl per well). For mGα proteins and β-arrestin recruitment assays [5], cells were transfected with Rluc8-mGαq or β-arrestin2-RlucII (BRET donors), rGFP-CAAX or tdrGFP-Rab5a (BRET acceptors), FLAG-$PAR_2$-HA and 3HA-$H_1$R. For GEMTA assays [6], cells were transfected with p63-RhoGEF-RlucII (BRET donor), rGFP-CAAX or tdrGFP-Rab5a (BRET acceptors), Gαq, FLAG-$PAR_2$-HA,3HA-$H_1$R, DynK44A and β-arrestin2. At 48 h post-transfection, cells were washed with PBS and the medium was replaced with HBSS containing 10 mM HEPES (pH 7.4). Cells were stimulated with trypsin (Sigma-Aldrich; #T0303) or histamine dihydrochloride (Tocris Bioscience; #354550) for 15 min, washed and then incubated histamine or trypsin, respectively, for 15 min before BRET measurement. The luciferase substrate Prolume purple (2 µM; Nanolight Technology; #369) was added for 6 min before BRET measurements. BRET was measured using a Synergy Neo2 Microplate reader (BioTek) with BRET2 filters (Donor filter: 410nm; acceptor filter: 515nm). The BRET signal was calculated as the ratio of light emitted by the energy acceptor over the light emitted by the energy donor.

**Statistics.** All results were analyzed and graphed using GraphPad Prism 10.2.3. Data were tested for normal distribution using the Prism package test for normal distribution including D'Agostino-Pearson omnibus normality test. For the analysis of the afferent nerve recordings, basal frequencies were first tested for outliers and for normality using the D'Agostino-Pearson omnibus normality test. They were then analyzed using a Wilcoxon paired test or the paired T-test. The amplitude of response to distensions was analyzed using 2-way RM ANOVAs with Šidák's *post hoc* test. For patch clamp experiments, data were analyzed using Welch's t test, or 1- or 2-way ANOVAs with Tukey's or Dunnett's test or, when indicated, non-parametric Kruskal-Wallis with Dunn's *post hoc* test. In afferent nerve recordings the number of mice used is represented by N, with one mouse being used per afferent nerve recording. The number of patient samples used was represented by n. The number of neuronal units present was represented by U. ebBRET data were analyzed using 2-way ANOVA followed by Dunnett's or Šídák's test. Results are expressed as mean±SEM with significance $P$<0.05.

### Figures

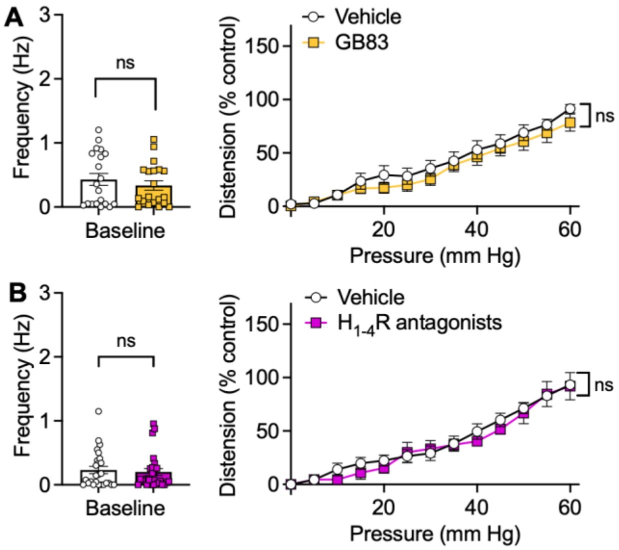


**Figure S1. Activity of colonic afferent neurons after intraluminal administration of vehicle.** Baseline activity and distension-induced responses of colonic afferent neurons after intraluminal administration of vehicle (control). Tissues were pre-treated with vehicle (control) or $PAR_2$ antagonist (GB83, 10 µM) (**A**) or $H_{1\text{-}4}R$ antagonists (1 µM pyrilamine, 10 µM ranitidine, 30 nM clobenpropit, 1 µM JNJ7777120) (**B**). Mean±SEM, N=number of mice, U=number of neuronal units. A: N=13, U=20. B: N=13 U=26. 2-way ANOVA or Wilcoxon paired test.

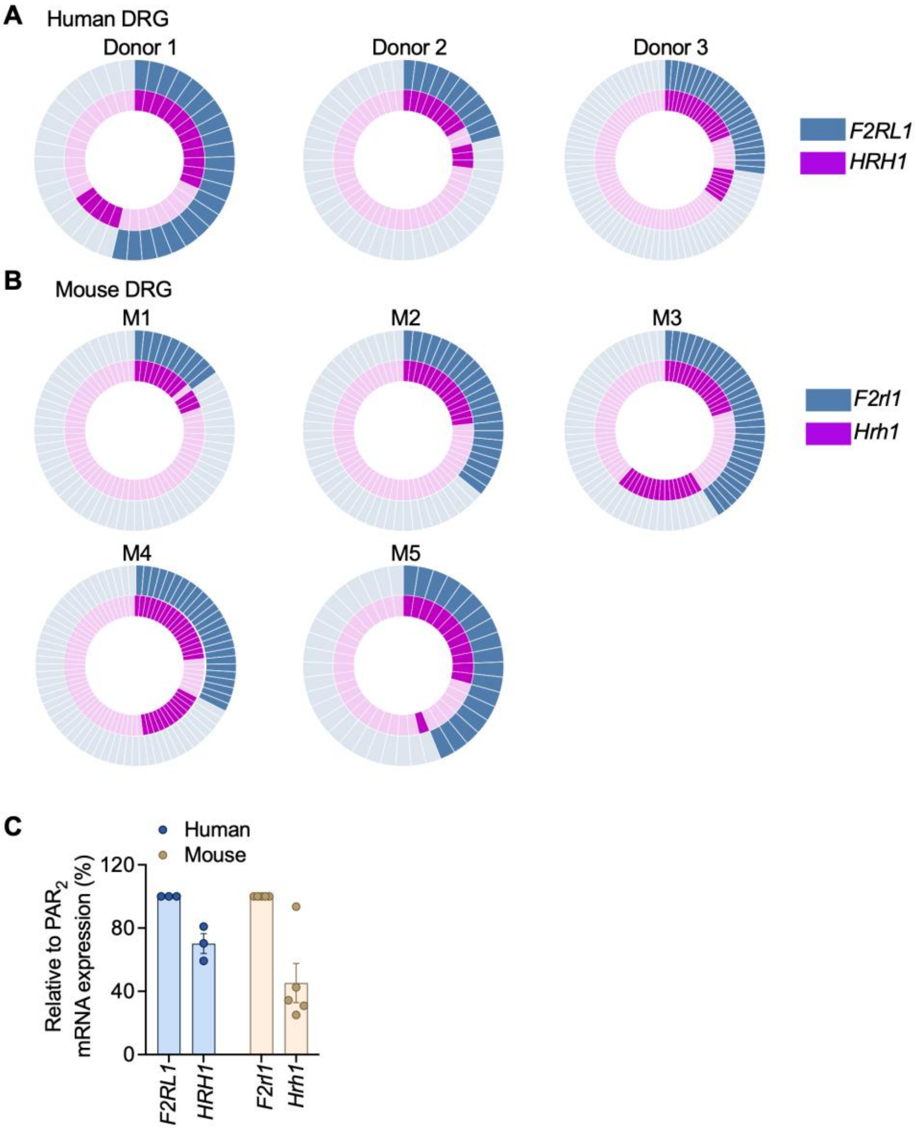


**Figure S2. Localization of $PAR_2$ and $H_1R$ mRNA in human and mouse DRG by RNAscope *in situ* hybridization.** Representative diagrams showing the total number of cells analyzed expressing *F2RL1/F2rl1* ($PAR_2$), *HNRH1/Hrh1* ($H_1R$) or both receptors in human DRG neurons (**A**) and mouse DRG neurons colocalizing with NeuN immunoreactivity (**B**). **C.** Percentage of cells co-expressing *F2RL1/F2rl1* and *HNRH1/Hrh1* relative to positive *F2RL1/F2rl1* DRG neurons. Data from n=3 humans and n=5 mice per group.

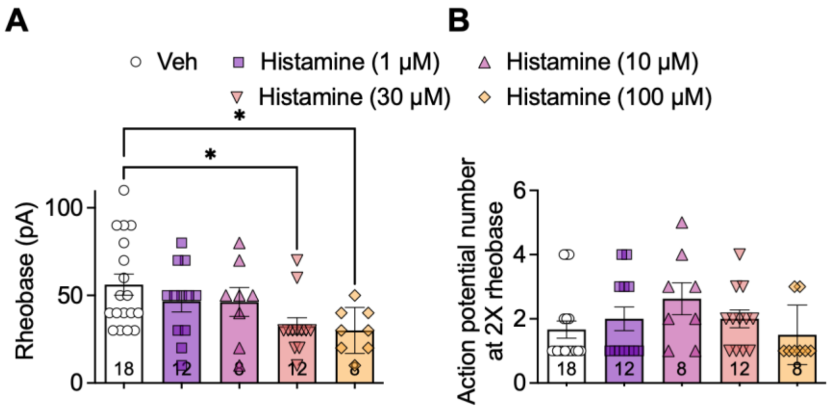


**Figure S3. Histamine concentration rheobase-response curve.** Pooled data showing rheobase (**A**) and the number of action potentials at twice rheobase (**B**) for mouse DRG neurons treated with vehicle or with histamine (1, 10, 30, 100 μM). Mean±SEM, data point indicate the number of neurons from N=6-8 mice (mean±SEM). *$P$< 0.05, Kruskal-Wallis with Dunn's *post hoc* test.

### Tables

#### Table S1. IBS-D patient pain score information

| PATIENT | SEX | PROMIS | IBS-SSS | | | | | |
|---|---|---|---|---|---|---|---|---|
| | | | Abdominal pain severity | Abdominal pain frequency | Abdominal distension | Bowel habits | Interference with daily life | Total SSS |
| **P1** | F | 23 | 75 | 75 | 75 | 90 | 75 | 390 |
| **P2** | F | 25 | 77 | 81 | 95 | 96 | 98 | 447 |
| **P3** | F | 25 | 55 | 40 | 38 | 99 | 95 | 327 |

#### Table S2. IBS-SSS Survey Questions

| | | | |
|---|---|---|---|
| 1 | Do you currently suffer from abdominal pain? | YES/NO | 0_____________100 |
| 2 | Enter the number of days you get pain in every 10 days | Numbers of days with pain_____x10 | |
| 3 | Do you currently suffer from abdominal distention? | YES/NO | 0_____________100 |
| 4 | Indicate how satisfied you are with your bowel habits | 0_____________100 | |
| 5 | Indicate how much IBS affects or interferes with your life in general | 0_____________100 | |

#### Table S3. PROMIS belly pain score survey questions

| | | |
|---|---|---|
| 1 | In the past 7 days how often did you have belly pain? | 1 → 5 |
| 2 | In the past 7 days at its worst how would you rate your belly pain? | 1 → 5 |
| 3 | In the past 7 days how much did your bely pain interfere with your day-today activities? | 1 → 5 |
| 4 | In the past 7 days how much did belly pain bother you? | 1 → 5 |
| 5 | In the past 7 days how often did you have discomfort in your belly? | 1 → 5 |

**Table S4. Lumbar DRG donor information for RNAScope analysis.** *DNC, death by neurological criteria.

| Donor number | Age | Sex | Race | Cause of death |
|---|---|---|---|---|
| 1 | 36 | M | White | Anoxia/Asphyxiation |
| 2 | 58 | M | White | DNC* |
| 3 | 47 | M | White | DNC* |
